# Transmission dynamics and between-species interactions of multidrug-resistant Enterobacteriaceae

**DOI:** 10.1101/436006

**Authors:** Thomas Crellen, Paul Turner, Sreymom Pol, Stephen Baker, To Nguyen Thi Nguyen, Nicole Stoesser, Nicholas P.J. Day, Ben S. Cooper

## Abstract

Widespread resistance to antibiotics is among the gravest threats to modern medicine, and controlling the spread of multi-drug resistant Enterobacteriaceae has been given priority status by the World Health Organization. Interventions to reduce transmission within hospital wards may be informed by modifiable patient-level risk factors for becoming colonised, however understanding of factors that influence a patient’s risk of acquisition is limited. We analyse data from a one year prospective carriage study in a neonatal intensive care unit in Cambodia using Bayesian hierarchical models to estimate the daily probability of acquiring multi-drug resistant organisms, while accounting for patient-level time-varying covariates, including interactions between species, and interval-censoring of transmission events. We estimate the baseline daily probability for becoming colonised with third generation cephalosporin resistant (3GC-R) *Klebsiella pneumoniae* as 0.142 (95% credible interval [CrI] 0.066, 0.27), nearly ten times higher than the daily probability of acquiring 3GC-R *Escherichia coli* (0.016 [95% CrI 0.0038, 0.049]). Prior colonization with 3GC-R *K. pneumoniae* was associated with a greatly increased risk of a patient acquiring 3GC-R *E. coli* (odds ratio [OR] 6.4 [95% CrI 2.8, 20.9]). Breast feeding was associated with a reduced risk of colonization with both 3GC-R *K. pneumoniae* (OR 0.73 [95% CrI 0.38, 1.5]) and *E. coli* (OR 0.62 [95% CrI 0.28, 1.6]). The use of an oral probiotic (*Lactobacillus acidophilus*) did not show clear evidence of protection against colonization with either 3GC-R *K. pneumoniae* (OR 0.83 [95% CrI 0.51, 1.3]) or 3GC-R *E. coli* (OR 1.3 [95% CrI 0.77, 2.1]). Antibiotic consumption within the past 48 hours did not strongly influence the risk of acquiring 3GC-R *K. pneumoniae*. For 3GC-R *E. coli*, ceftriaxone showed the strongest effect for increasing the risk of acquisition (OR 2.2 [95% CrI 0.66, 6.2]) and imipenem was associated with a decreased risk (OR 0.31 [95% CrI 0.099, 0.76). Using 317 whole-genome assemblies of *K. pneumoniae*, we determined putatively related clusters and used a range of models to infer transmission rates. Model comparison strongly favored models with a time-varying force of infection term that increased in proportion with the number of colonized patients, providing evidence of patient-to-patient transmission, including among a cluster of *Klebsiella quasipneumoniae*. Our findings provide support for the hypothesis that *K. pneumoniae* can be spread person-to-person within ward settings. Subsequent horizontal gene transfer within patients from *K. pneumoniae* provides the most parsimonious explanation for the strong association between colonization with 3GC-R *K. pneumoniae* and acquisition of 3GC-R *E. coli*.

## 1 Introduction

Antimicrobial resistance (AMR) is among the gravest threats to modern medicine and presents a challenge to healthcare providers in all regions of the world [1]. Within hospitals Enterobacteriaceae such as *E. coli* and *Klebsiella pneumoniae* have been highlighted as presenting a particular threat given high levels of resistance to third generation cephalosporins, carbapenems and, in the case of *K. pneumoniae*, the ability to persist on surfaces and a plastic genome that can readily acquire and disseminate plasmids [2, 3]. Consequently these pathogens have been given priority status for control by the World Health Organization. Multidrug-resistant organisms (MDRO) can be imported into hospitals after exposure in the community, the environment, or in the case of neonates may be acquired congenitally during birth [4]. Onward transmission in healthcare settings may occur due to contact between patients, contamination of the environment, or via healthcare workers [5]. Carriage of potentially pathogenic Enterobacteriaceae in the gastrointestinal tract is assumed to be a prerequisite for bloodstream infection and invasive disease [6].

Facing the challenge of controlling MDRO will require evidence based interventions. Randomized controlled trials remain the gold-standard approach, but are costly and challenging to implement. Understanding the risk factors that impact on MDRO acquisition can inform interventions or control measures, and can be performed using routinely collected swab data. A common approach is to perform a case control study between carriers of, for instance, extended spectrum *β*-lactamase producing Enterobacteriaceae (ESBL-E) and either negative cases or carriers of non-ESBL-E [7]. Such approaches are problematic as they do not consider the process that gives rise to colonization, namely repeated exposure of the patient to sources of multidrug-resistant Enterobacteriaceae. Given a daily probability of colonization *p*, the cumulative probability of observing a colonization event increases each day the patient is in the ward, and on day *N* is given by:

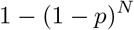

The common ‘finding’ that an increased length of stay is associated with carriage of ESBL-E [8, 9, 10] is therefore a consequence of recurrent exposure, and not matching cases and control by length of stay may produce spurious associations.

Another approach to studying the transmission of nosocomial pathogens has been through the use of compartmental models which modify the general epidemic model [11, 12, 13]. Within this framework, patients are generally assumed to be homogeneous with respect to acquiring or transmitting pathogens and the approach is not compatible with the inclusion of covariates. Further, as these models are rarely fit to data their relevance to clinical decision making can be limited [14]. An alternative to compartmental models are stochastic statistical models, which account for imperfect observation of the transmission process. As swabs are not taken every day, acquisition events are censored over the swabbing interval [15]. This statistical approach has considered the number of new colonizations in the ward within a swabbing interval as a binomial process, however this only permits the inclusion of covariates at the ward level [15]. As interventions to reduce the burden of AMR in hospitals are often directed at individual patients, such as the use of narrow spectrum antibiotics [16], shortening the duration of treatment [17] or administering probiotics [18], evaluating how these interventions influence the risk of an individual acquiring MDRO will be valuable in assessing their efficacy.

Here we develop novel statistical methods to, firstly, estimate the individual-level susceptibility to colonization per day with multi-drug resistant *K. pneumoniae* and *E. coli*, while incorporating time-varying risk factors for acquisition. We then develop models which allow for the incorporation of genomic or sequence-typing data to estimate transmission rates for separate clusters.

## 2 Methods

We estimated the daily probability of colonization with a MDRO under the assumption that individuals remain colonized for the duration of their stay until discharge [19]. Each day in the ward, a previously uncolonized patient can become colonized, however as we lack swabbing results from each day, the outcome is censored over a swabbing interval of *N* days. If the probability of becoming colonized on day *i* for patient *j* is *p_ij_*, given the patient is uncolonized at the start of the day, then the probability of remaining uncolonized is (1−*p_ij_*). In interval *k* for patient *j* consisting of *N_kj_* days, then the probability of remaining uncolonized is:

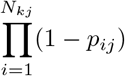

Therefore the probability of becoming colonized is:

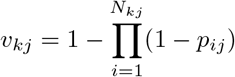

The outcome for patient *j* in interval *k*, *X*_*kj*_ ∈ {0,1}, as the patient either becomes colonized (1) or remains uncolonized (0) with a MDRO. Therefore the likelihood function is:

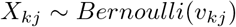

The daily probability of becoming colonized (*p_ij_*) is related by the logit link function to a linear function of covariates (*π_ij_*):

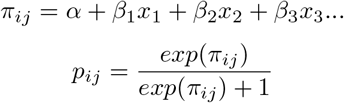

Where *x*_1_, *x*_2_, *x*_3_… is a vector of predictors (data) and *β*_1_, *β*_2_, *β*_3_… is a vector of parameters that are to be estimated. The intercept *α* can be a single parameter, or permitted to vary between period of time (months or weeks), or to vary by patient. The range of values *α* can take are assumed to be normally distributed, with a mean *μ* and variance *σ* which are themselves parameters with their own prior distributions.

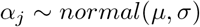

We used data collected prospectively from a neonatal unit (NU) in a children’s hospital in Siem Reap, Cambodia between September 2013 (when the NU newly opened) and September 2014. The study protocol required rectal swabs to be taken within 48 hours of admission and subsequently every three to four days. However in practice considerable variation was observed in the timing between swabs. We assumed that any patient testing positive within 48 hours of admission was positive on arrival, and therefore excluded from the analysis. Patients that were swabbed >48 hours from admission but were negative on their first swab were included in the analysis, while those that tested positive were omitted. Details of the microbiological treatment of rectal swabs, including resistance assays, have been published previously [18].

We developed and fitted models to examine risk factors for the acquisition of 3GC-R *K. pneumoniae* and, separately, *E. coli* in the NU for infants that were not colonized by the respective 3GC-R organism on admission. We examined a range of models, alternating i) single intercepts, and models where the intercept was permitted to vary by random effects (weeks, months or patients), ii) the inclusion of interaction terms between commonly used pairs of antibiotics and iii) the inclusion of a force of infection term for the pseudo mass-action principal [15].

We included as covariates exposure to the five most commonly taken antibiotics (ampicillin, ceftriaxone, imipenem, gentamicin and cloxacillin) within the past 48 hours, whether the infant was breastfed, if the infant recieved an oral probiotic on entry (*Lactobacillus acidophilus*), sex, prematurity, severity (defined as either i) requiring ventilation, ii) requiring continuous positive airway pressure or iii) prolonged rupture of membranes), and if already colonized with 3GC-R *E.coli* in the case of 3GC-R *K. pneumoniae* acquisition and vice versa. These explanatory variables were treated as binary (0/1). We also included the age in days on first admission to the NU, which was treated as continuous. Covariates were recorded for every day the infant was present in the NU.

We whole-genome sequenced 317 cultured isolates identified morphologically as *K. pneumoniae* from i) rectal swabs from all colonized patients within a four month period of the study and ii) twice weekly swabs from 7 environmental surfaces around the ward (6 sinks and 1 computer keyboard) within the same time frame. Sequencing was performed with the Illumina HiSeq 2500 platform, producing 150 base-pair paired-end reads. The reads were trimmed for adapter sequence using TrimGalore v0.4.4 before assembly with Unicycler v0.4.5 [20], contigs <1 kilobase were discarded. Distances between assemblies were calculated using mash v1.1 [21] and a phylogeny constructed with mashtree v0.33. Cluster cut-offs were defined for mash distances of >0.0175 and clusters are considered as groups of putatively related isolates. Sequence types (STs) were identified using Kleborate 0.2.0 [22].

We assessed within-ward transmission of 3GC-R *K. pneumoniae* clusters based on three transmission models, 1) where colonization with an isolate from cluster *c* is independent of the number of individuals colonized with cluster *c*, 2) including a coefficient for the force of infection, where the covariate is the number of patients colonized with cluster *c* in the NU on day *i*, 3) as model 2 but without an intercept. The probability of colonization for individual *j* on day *i* with cluster *c* for the respective models are:

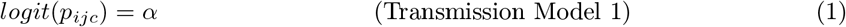

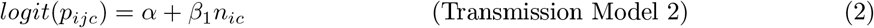

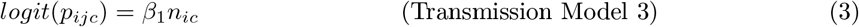

For the transmission models, the probability of colonization within a swabbing interval of *N_kj_* days is calculated as shown above, and the interval probability is used within the Bernoulli likelihood function.

We fitted the statistical models using Hamiltonian Markov Chain Monte Carlo in Stan (version 2.17.3) within the R environment (v. 3.4.3). Prior distributions were normal distributions using vaguely informative priors according to the principal of maximum entropy [23]. Chains were run for a varying number of iterations depending on the number of parameters to estimate, though with a minimum of 20,000 iterations including burn-in. The Gelman-Rubin statistic was used as a diagnostic (values>1 indicate incomplete convergence), additionally posterior chains were visually inspected for convergence. Model comparison was performed with widely applicable information criterion (WAIC) and leave-one-out cross validation (LOO-CV). We use 95% credible intervals (CrIs) as a measure of uncertainly around posterior parameter distributions.

## 3 Results

Over the year long observation of the cohort, high rates of carriage with 3GC-R-E were observed among the 333 infants admitted. For 3GC-R *K. pneumoniae*, 121 infants were colonized on first admission (first positive swab within 48 hours), and 21 were positive on their first swab taken after 48 hours and omitted. For the 191 infants susceptible on first admission, swab results were censored over 421 intervals, although for 19 of these no final outcome was recorded leaving 402 swabbing intervals consisting of 871 patient days in our analysis. Overall, 109/191 susceptible infants acquired 3GC-R *K. pneumoniae* during their stay in the NU. For 3GC-R *E. coli*, 97 patients were colonized on admission and additionally 14 were positive on their first swab taken after 48 hours and omitted. For the 222 susceptible infants, swab results were censored over 721 intervals, although the final outcome was missing for 32 of these, leaving 689 swabbing intervals consisting of 1728 patient days for inclusion in the analysis. Overall, 77/222 susceptible infants acquired 3GC-R *E. coli* during their stay in the NU. Among the cultured 3GC-R *K. pneumoniae* and 3GC-R *E. coli*, extended spectrum *β* lactamases were detected in 98.5% and 96.6% of samples respectively. Colonization with 3GC-R *Acinetobacter baumannii* and *Pseudomonas aeruginosa* was also observed, though the number of acquisition events (5 and 1 respectively) were insufficient for statistical analysis. Swabbing intervals were a median of 2 days (range 1, 10 days). Further details on the study population, including blood-stream infections and mortality have been reported previously [18], and a summary is provided in Table 1.

**Table 1:** Summary of cohort patient characteristics. 1. Interquartile range, 2. Colonized at entry is defined as an initial positive swab within the first 48 hours of admission, if the first swab is positive and it was taken later than 48 hours from admission then the infant is considered to be colonized at an unknown time, 3. Assigned by clinician to receive oral *Lactobacillus acidophilus*, 4. Severe symptoms are requiring ventilation, continuous airway pressure or prolonged rupture of membranes.

| Variable | Males | Females | Total |
| --- | --- | --- | --- |
| Number of Patients | 177 (53.1%) | 156 (47.8%) | 333 |
| Length of Stay in Days<br>Median, (IQR)<sup>1</sup> | 6 (4, 11) | 5 (4, 9) | 6 (4, 10) |
| Colonised with<br>*K. pneumoniae* at Entry<br>(or Unknown Time)<sup>2</sup> | 66 (10) | 55 (11) | 121 (21) |
| Colonised with<br>*K. pneumoniae*<br>During Admission | 54/101 (54.5%) | 55/90 (61.1%) | 109/191 (57.1%) |
| Colonised with *E. coli*<br>at Entry<br>(or Unknown Time)<sup>2</sup> | 53 (10) | 44 (4) | 97 (14) |
| Colonised with *E. coli*<br>During Admission | 38/114 (33.3%) | 39/108 (36.1%) | 77/222 (34.7%) |
| Age at Admission in Days<br>(IQR)<sup>1</sup> | 8 (2, 15) | 9 (1, 17) | 8 (1, 16) |
| Probiotic Taken<sup>3</sup> | 76/177 (42.9%) | 62/156 (39.7%) | 138/333 (41.4) |
| Breast Milk Fed | 163/177 (92.1%) | 139/156 (89.1%) | 302/333 (90.1%) |
| Severe<sup>4</sup> | 35/177 (19.8%) | 32/156 (20.5%) | 67/333 (20.1%) |
| Born Premature | 30/177 (16.9%) | 24/156 (15.4%) | 54/333 (16.2%) |

The temporal trend in counts of infants colonized per day are shown in Figure 1 for both 3GC-R *K. pneumoniae* and 3GC-R *E. coli* (panels A and B), with imported cases considered separately to cases acquired in the ward. A representation of the swabbing interval outcomes for the first thirty susceptible patients for the respective organisms are shown in panels C and D.

**Figure 1:**
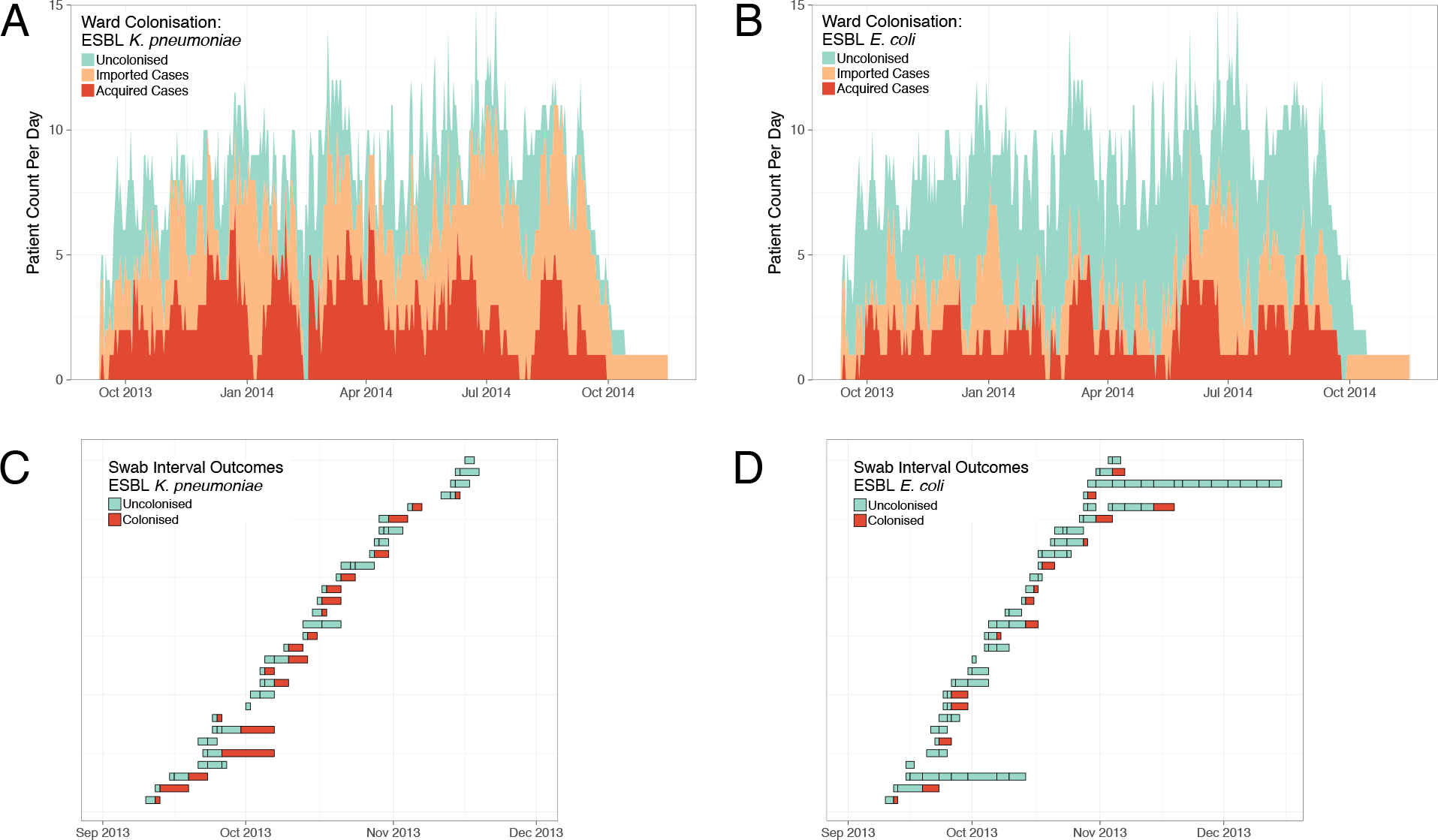
Descriptive Epidemiological Data of a Cohort of Neonates Colonized with Third Generation Cephalosporin Resistant (3GC-R) Bacteria from a Children’s Hospital in Cambodia. Daily counts of neonates colonized with 3GC-R *Klebsiella pneumoniae* and *Escherichia coli* are shown in panels A and B respectively over the year long study period, where colour reflects uncolonized, imported or acquired cases. The total height of the bars shows the ward occupancy on that day. The results from rectal swabs among the first thirty patients uncolonized at entry for 3GC-R *Klebsiella pneumoniae* and *Escherichia coli* are shown in panels C and D respectively. Each row represents a patient and each colored block represents a swab interval, the width of the interval is the number of days in that interval. Outcomes are shown up to the first positive swab, after which time the patient is assumed to be colonized until discharge.

A number of models were fitted to *K. pneumoniae* and *E. coli* swab interval data. The simplest single-intercept model (1) was found to outperform models that included interaction terms for pairs of commonly used antibiotics (2) or a force of infection term (3) on the basis of WAIC and LOO-CV. Similarly, the simplest model for *E. coli* acquisition (4) outperformed models with the additional terms (5, 6). We therefore present the results from models 1 and 4.

The model estimates the daily baseline probability for a neonate becoming newly colonization with 3GC-R *K. pneumoniae* as 0.14 (95% CrI 0.066, 0.27) and with 3GC-R *E. coli* as 0.016 (95% CrI 0.0038, 0.049). The largest risk factor that modified the probability of acquisition with 3GC-R *K. pneumoniae* is prior colonization with 3GC-R *E. coli* (OR 2.1 [95% CrI 1.2, 3.6]), while the largest risk factor for acquisition of 3GC-R *E. coli* was prior colonization with 3GC-R *K. pneumoniae* (OR 6.4 [95% CrI 2.8, 20.9]). Male sex was the strongest protective effect against acquisition of 3GC-R *K. pneumoniae* (OR 0.66 [95% CrI 0.43, 1.0]), followed by breast milk feeding (OR 0.73 [95% CrI 0.38, 1.5]). The covariate with the strongest protective effect against 3GC-R *E. coli* was the use of imipenem within the past 48 hours (OR 0.31 [95% CrI 0.099, 0.76]), again followed by breast milk feeding (OR 0.62 [95% CrI 0.28, 1.6]). The use of an oral probiotic, which was administered at the discretion of the administering clinician, did not show a strong effect on either the acquisition of 3GC-R *K. pneumoniae* (OR 0.83 [95% 0.51, 1.3]), or on 3GC-R *E. coli* (OR 1.3 [95 % CrI 0.77, 2.1]). The effect of consuming any of the five antibiotics over the past 48 hours did not substantially alter the daily risk of acquiring 3GC-R *K. pneumoniae* (80% CrIs all overlap unity; see Figure 2 panel A). For the acquisition of 3GC-R *E. coli*, consumption of ceftriaxone (OR 2.2 [95% CI 0.66, 6.2]) and cloxacillin (OR 1.7 [0.71, 4.0]) in the past 48 hours increased the risk of becoming colonized. Ampicillin and gentamicin showed, respectively, a small protective and deleterious effect for colonization with 3GC-R *E. coli*, however as these antibiotics were commonly administered together and the posterior distributions of both are centered around unity, this suggests no correlation. The carbapenem imipenem, which has action against ESBL-producing organisms, had a protective effect as described above. Infants that qualified as severe (see Methods) did not show an increased risk of acquisition for 3GC-R *K. pneumoniae* (1.1 [95% CI 0.58, 2.2]) or *E. coli* (0.96 [95% 0.55, 1.7]). A premature birth increased an infant’s risk of becoming colonized with 3GC-R *K. pneumoniae* by an OR of 1.8 (95 % CI 0.96, 3.5), though the association with acquisition of 3GC-R *E. coli* was weaker and in the opposite direction (OR 0.84 [95% CI 0.44, 1.6]). Posterior predictions for the impact of covariates on the probability of colonization are shown in Tables 2 and 3, and Figure 2 (panels A and B).

**Figure 2:**
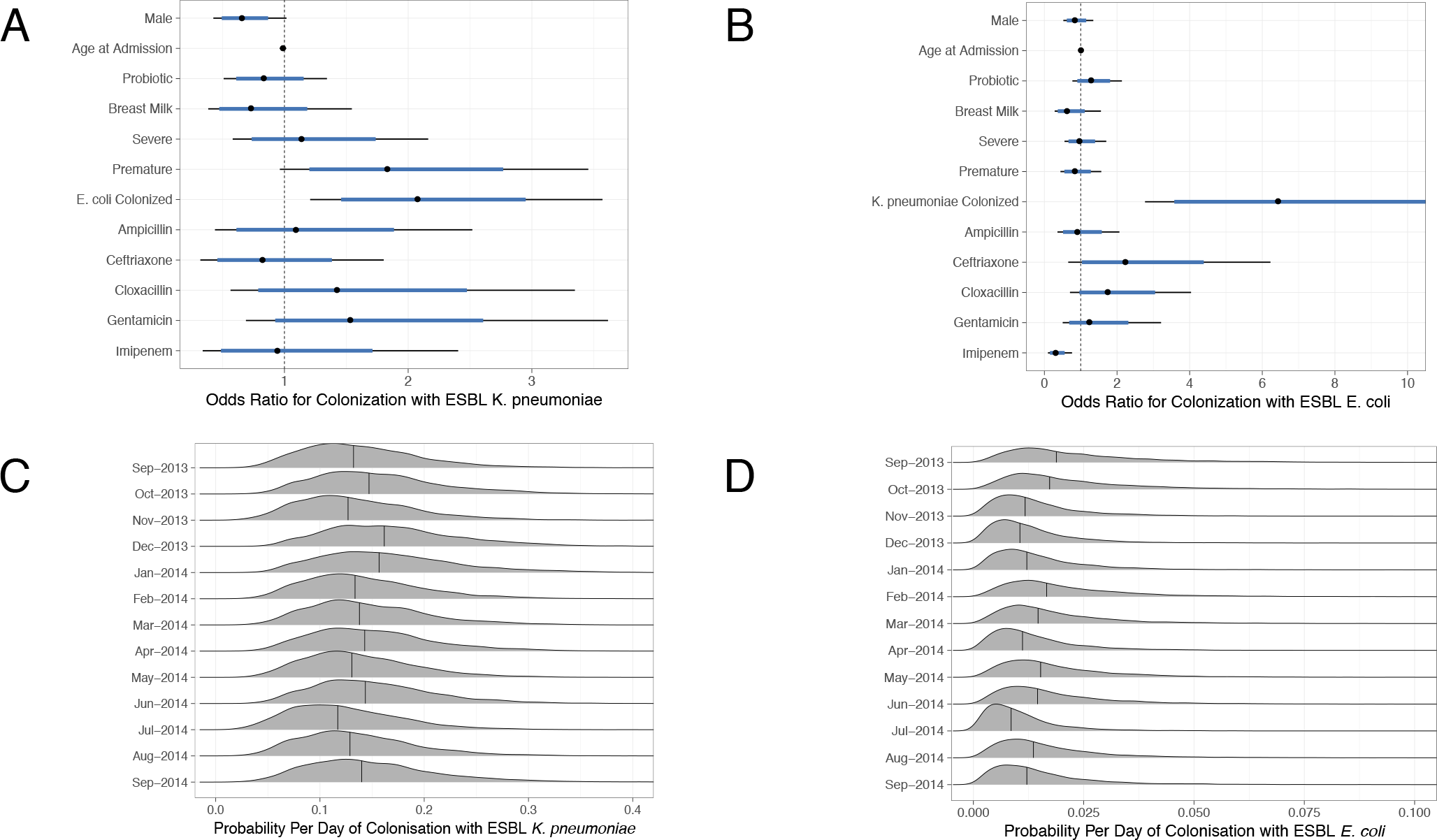
Covariates and Intercepts for Susceptibility to Colonization. The posterior distribution of odds ratio for covariates on the daily risk of becoming colonized with third generation cephalosporin resistant (3GC-R) *Klebsiella pneumoniae* and *Escherichia coli* are shown in panels A and B respectively. The thick blue line represents the 80% credible interval (CrI) and the the thin black line represents the 95% credible interval. Note that panel B is truncated as the 95% CrI odds ratio for acquiring 3GC-R *E. coli* extends to 20 (see Table 3). The posterior distributions of the intercepts (baseline risk) for acquisition of *K. pneumoniae* and *E. coli* are shown by month in panels A and B respectively, taken from a hierarchical model where the intercept was permitted to vary by study month.

**Table 2:** Risk factors and the daily probability for acquisition of third-generation cephalosporin resistant (3GC-R) *Klebsiella pneumoniae* among neonates admitted to an intensive care unit in Angkor Hospital for Children, Cambodia between September 2013 - September 2014. Posterior parameter distributions are shown on the odds ratio scale and the probability scale. 1. Credible Interval of Posterior Distribution, 2. Probability of acquisition is the posterior intercept plus the posterior coefficients transformed onto the probability scale, 3. Assigned by clinician to receive oral *Lactobacillus acidophilus*, 4. Severe symptoms are requiring ventilation, continuous airway pressure or prolonged rupture of membranes, 5. Patient is already colonised with 3GC-R *Escherichia coli*, 6. Antibiotics consumed within the past 48 hours.

| Variable | Odds Ratio<br>Posterior Median<br>(95% CrI<sup>1</sup>) | Daily Probability<br>of Acquisition<sup>2</sup><br>Posterior Median<br>(95% CrI<sup>1</sup>) |
| --- | --- | --- |
| Baseline | Reference | 0.14 (0.066, 0.27) |
| Male | 0.66 (0.43, 1.0) | 0.098 (0.043, 0.20) |
| Age at Admission<br>(Days) | 0.99 (0.96, 1.0) | 0.14 (0.066, 0.26) |
| Probiotic Taken<sup>3</sup> | 0.83 (0.51, 1.3) | 0.12 (0.054, 0.24) |
| Breast Milk Fed | 0.73 (0.38, 1.5) | 0.11 (0.058, 0.18) |
| Severe<sup>4</sup> | 1.1 (0.58, 2.2) | 0.16 (0.062, 0.33) |
| Born Premature | 1.8 (0.96, 3.5) | 0.23 (0.099, 0.44) |
| *E. coli*<br>Colonised<sup>5</sup> | 2.1 (1.2, 3.6) | 0.26 (0.11, 0.47) |
| Ampicillin<sup>6</sup> | 1.1 (0.44, 2.5) | 0.15 (0.056, 0.33) |
| Ceftriaxone<sup>6</sup> | 0.82 (0.32, 1.8) | 0.12 (0.037, 0.29) |
| Cloxacillin<sup>6</sup> | 1.4 (0.56, 3.3) | 0.19 (0.071, 0.39) |
| Gentamicin<sup>6</sup> | 1.5 (0.69, 3.6) | 0.20 (0.077, 0.44) |
| Imipenem<sup>6</sup> | 0.94 (0.34, 2.4) | 0.13 (0.043, 0.32) |

**Table 3:**
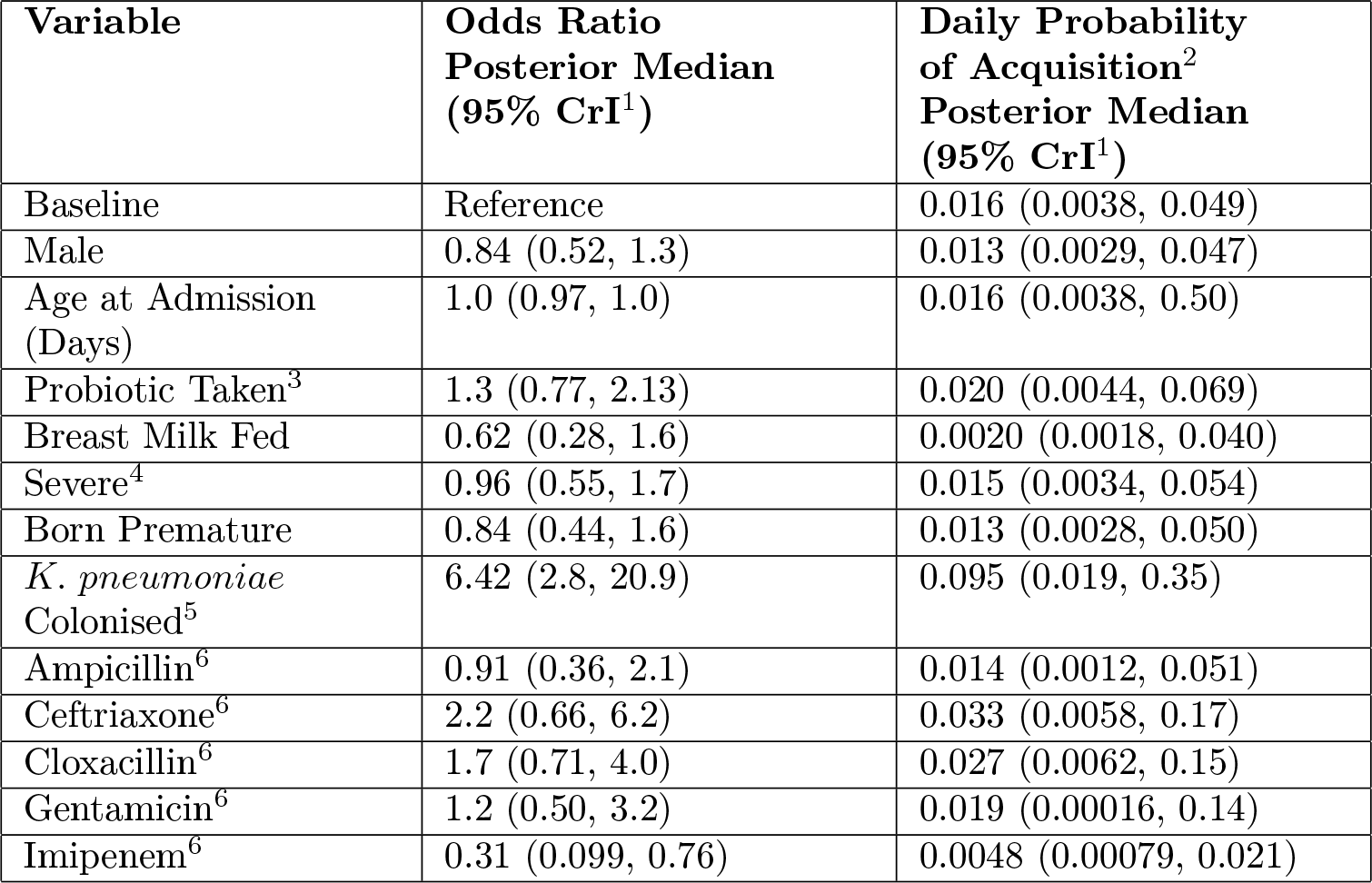
Risk factors and the daily probability for acquisition of third-generation cephalosporin resistant (3GC-R) *Escherichia coli* among neonates admitted to an intensive care unit in Angkor Hospital for Children, Cambodia between September 2013 - September 2014. Posterior parameter distributions are shown on the odds ratio scale and the probability scale. 1. Credible Interval of Posterior Distribution, 2. Probability of acquisition is the posterior intercept plus the posterior coefficients transformed onto the probability scale, 3. Assigned by clinician to receive oral *Lactobacillus acidophilus*, 4. Severe symptoms are requiring ventilation, continuous airway pressure or prolonged rupture of membranes, 5. Patient is already colonised with 3GC-R *Klebsiella pneumoniae*, 6. Antibiotics consumed within the past 48 hours.

In models where we included the force of infection (a parameter linked the number of other colonized patients in the NU on the same day), this was found to have a negative slope for both acquisition of 3GC-R *K. pneumoniae* and *E. coli*. Epidemiological theory suggests that if patient-to-patient transmission was occurring, we would expect the force of infection for any organism to increase with the number of individuals colonized in the ward with that organism; therefore the negative slope we observe in our models suggests that the force of infection is non-identifiable either i) as a result of the high counts of colonized patients at almost every time point (Figure 1) or ii) due to structure in the pathogen population.

We also investigated models where the intercept (probability of colonization in the absence of covariates), was permitted to vary over periods of time (weeks, months) or by individual patients. High temporal or individual-level variation in intercepts would indicate that transmission is due to sporadic, time-limited outbreaks. Due to the small number of observations per cluster, we generally observed low variation between weekly or per-patient clusters as the estimates were shrunk towards the global mean. In a model where the intercept was permitted to vary by month, we observed little monthly variation for both organisms suggesting that the force of infection remained relatively constant over the observed period (Figure 2 panels C and D).

To better understand the structure of the pathogen population, we examined whole-genome assemblies of 317 3GC-R *K. pneumoniae* isolates taken from rectal or environmental swabs in the NU over four months. A phylogeny based on k-mer distances between assembles is shown in Figure 3 (panel A), of note is the highly diverse and structured nature of the pathogen population, in contrast to one dominated by a clonal expansion of a single lineage. Overall 62 distinct STs were observed in our collection of isolates. The species identified morphologically from culture as *Klebsiella pneumoniae* consists of three distinct subpopulations that meet the criteria for separate species, these are defined in the literature as *K. pneumoniae* (KpI), *K. quasipneumoniae* (KpII) and *K. variicola* (KpIII). We have isolated all three species from infants in the cohort (KpI, *n*= 219, KpII, *n*= 95, KpIII, *n*= 3), and furthermore the diversity is markedly similarly to a global collection of *K. pneumoniae* isolates [24], suggesting that the diversity observed within a Cambodian NU over four months is comparable to the diversity of *K. pneumoniae* globally.

**Figure 3:**
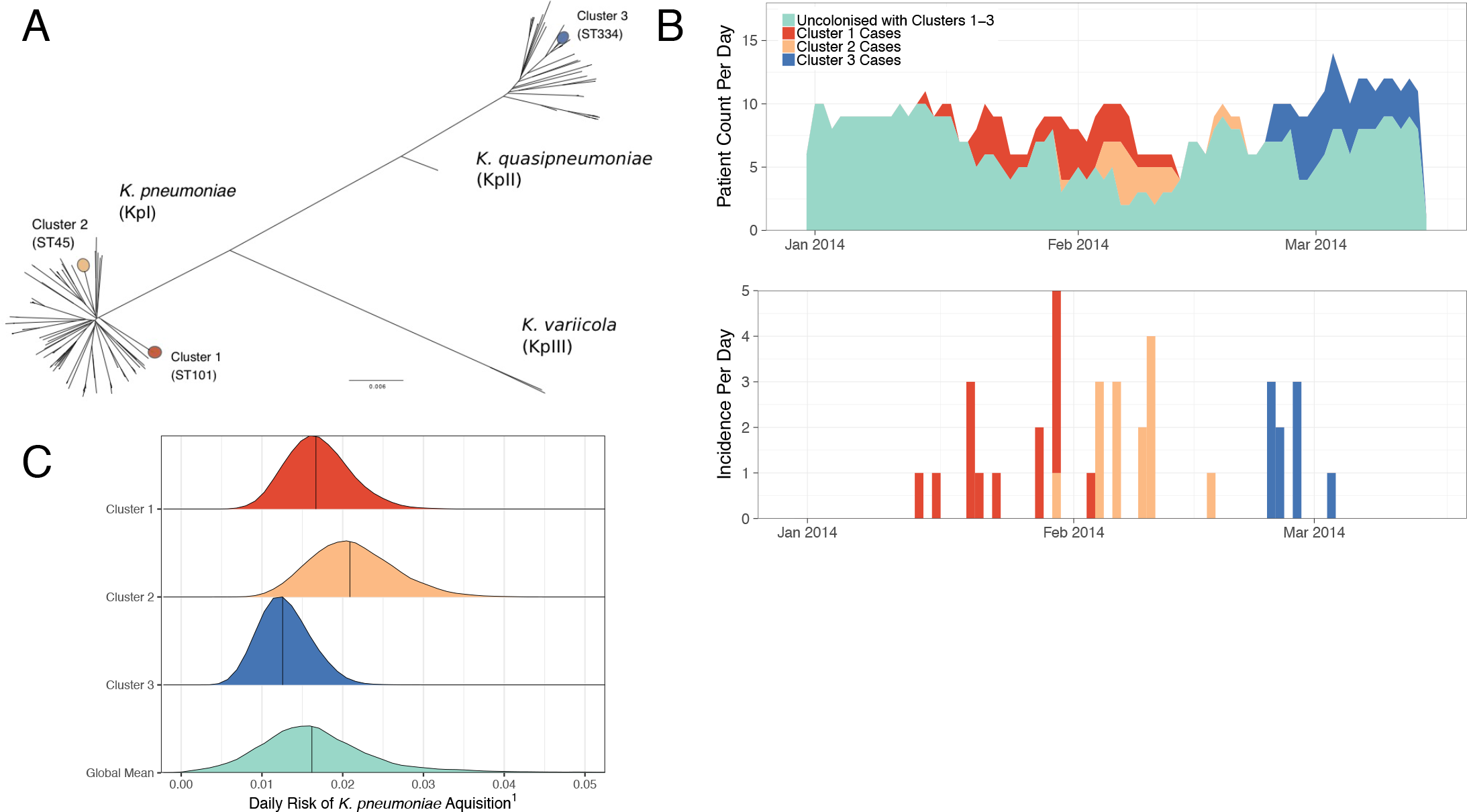
Phylogeny of third generation cephalosporin resistant (3GC-R) *Klebsiella pneumoniae* collected from rectal swabs and the environment from a neonatal care unit in Cambodia. The unrooted phylogeny of 317 whole genome assemblies is shown in panel A, where the branch length gives the mash distance (a measure of k-mer similarity) between assemblies. The three largest clusters used in the transmission analysis are highlighted on the tree. Note that the species identified morphologically as *K. pneumoniae* is a complex consisting of three distinct clusters that are considered to be separate species (*K. pneumoniae*, *K. quasipneumoniae* and *K. variicola*). The number of cases per day and the incidence of colonization with clusters 1-3 are shown in panel B over the period January to March 2014. The daily risk of colonization with *K. pneumoniae* from clusters 1-3 are shown in panel C as a density plot of the posterior distribution from transmission model 2. Note that the x-axis (1) denotes the risk of becoming colonized for any susceptible infant, per colonized patient in the ward, per day. When this is multiplied by the number of colonized patients in the ward it gives the force of infection.

In order to estimate transmission parameters for clusters of *K. pneumoniae*, we first defined 82 putatively related clusters using a distance cut-off based on partitions in the phylogeny of which the median cluster size is 2 isolates (range 1, 25). Investigating the largest of these, cluster 1 (within KpI) contains 25 isolates that are all ST101 (21 from patient swabs, 4 from environmental swabs), cluster 2 (KpI) contains 19 isolates that are all ST45 (17 from patient swabs, 2 from environmental swabs) and cluster 3 (KpII) contains 16 isolates that are all ST334 (14 from patient swabs and 2 from environmental swabs). Figure 3 shows the location of these clusters on the phylogeny (panel A) and the temporal incidence of cases from each cluster (panel B). Cases within each of the three clusters occurred over relatively short periods of 20, 18, and 10 days respectively, suggesting that the cases may be linked epidemiologically either through i) patient-to-patient transmission (via the hands of healthcare workers) or ii) from a common source outbreak.

We fitted a range of models to acquisition of the three clusters as described in the Methods. Model comparison showed that most weight was assigned to model 2, supporting the inclusion of the force of infection parameter and indicating that patient-to-patient transmission was occurring for these three clusters. Model 2 was re-fit as a hierarchical model where the force of infection (*β*_1_) was permitted to vary by cluster (with priors for the global mean and variance). The results are shown in Figure 3 (panel C).

Overall the three clusters have a similar effect on raising the risk of acquisition, with posterior medians of 0.017, 0.021 and 0.012 respectively. The cluster 3 isolates are from *Klebsiella quasipneumoniae* (Figure 2, panel A), which is considered to be less pathogenic than *K. pneumoniae* [24], however our results suggest that this organism can be spread person-to-person within health-care settings.

## 4 Discussion

We have inferred the transmission dynamics of multi-drug resistant Enterobacteriaecae using novel statistical methods. Our results indicate that prior colonization with another 3GC-R-producing organism is the largest risk factor for acquisition of 3GC-R *K. pneumoniae* and 3GC-R *E. coli*, with the strongest effect in the *K. pneumoniae* → *E. coli* direction (OR 6.4, see Table 3 and Figure 2 panel B). Plasmid-mediated spread of 3GC-R producing genes between these two organisms has been documented previously among neonates [25], and a number of whole-genome sequencing studies have demonstrated the spread of carbapenemase producing genes on plasmids by *K. pneumoniae* [26, 27]. Therefore reducing the spread of *K. pneumoniae* in particular will have a disproportionate effect on the overall prevalence of 3GC-R carriage.

Modeling the acquisition of *K. pneumoniae* within related clusters permitted the estimation of the force of infection, which was not possible in models that did not consider clusters separately. Comparison between competing models supported the inclusion of a force of infection term and provides strong evidence that patient-to-patient transmission in the NU, mediated by healthcare workers, is occurring. We observed some variation in the probability of per-day colonization with the three clusters (Figure 3 panel C), and cluster 3, which consists of *K. quasipneumoniae*, had a lower force of infection, which may indicate lower transmissibility. As we selected the largest clusters to estimate the force of infection, the values we obtained may be biased upwards.

Breast milk feeding was associated with a reduced risk of colonization with both 3GC-R *K. pneumoniae* and *E. coli*. This corresponds with our understanding of the development of a healthy gut microbiome in the early stages of life, which can be adversely affected by the replacement of breast milk by formula [28], and the protective effect of a diverse microbiome that competes against potentially pathogenic bacteria [29]. The finding that the oral probiotic *Lactobacillus acidophilus* was not protective against colonization with either organism was disappointing in light of earlier results that appeared to show an effect [18], however our result corresponds with recent findings that probiotics given after antibiotic consumption impairs and delays the recovery of normal gut flora in humans [30].

All but one class of antibiotics was associated with an increased risk of acquisition of 3GC-R *E. coli*, in particular the third-generation cephalosporin ceftriaxone (OR 2.2 [95% CrI 0.66, 6.2]). The exception to this was imipenem (OR 0.31 [95% CrI 0.099, 0.76]), which exhibited a protective effect as carbapenems are the last line against 3GC-R producing organisms and a low level or carbapenamase resistance was observed in this cohort. The effect of antibiotic consumption on 3GC-R *K. pneumoniae* acquisition was more mixed, with most drugs exhibiting modest effect sizes and the 80% CrIs overlapped unity in all cases (Figure 2, panel A).

Our study has a number of limitations. We assumed that a culture positive patient within 48 hours of admission is colonized at entry, along with the diagnostic being 100% sensitive and specific. The models we presented could be extended to account for the uncertainty in the diagnostic process, although this would increase model complexity and the computational intensity of model fitting. We discussed risk factors for the acquisition of two organisms, however we sequenced isolates only of *K. pneumoniae* from a four month period; a more complete dataset would also have sequenced *E. coli* and included long read sequence data to reliably infer plasmid transmission [26].

By developing models that can be fitted to routinely observed data we have developed a powerful framework to assess the transmission dynamics of multidrug resistant Enterobacteriaceae and perform parameter inference for risk factors for infants becoming colonized and a force of infection term. Furthermore we have used colonization data from one of the largest carriage studies to provide insight in a clinical setting with a high prevalence of MDROs, where the need to intervene is high. Future work will estimate the impact of potential interventions by developing agent-based models and simulating from the parameter estimates we have obtained in this study.

## References

[1] World Health Organization. Antimicrobial resistance: global report on surveillance. World Health Organization, 2014.

[2] Anton Y. Peleg and David C. Hooper. Hospital-acquired infections due to gram-negative bacteria. New England Journal of Medicine, 362(19):1804–1813, 2010.

[3] Kelly L. Wyres and Kathryn E. Holt. Klebsiella pneumoniae population genomics and antimicrobial-resistant clones. Trends in microbiology, 24(12):944–956, 2016.

[4] A. K. M. Zaidi, W. C. Huskins, D. Thaver, Z. A. Bhutta, Z. Abbas, and D. A. Goldmann. Hospital-acquired neonatal infections in developing countries. The Lancet, 365(9465):1175–1188, 2005.

[5] Donald A Goldmann. Bacterial colonization and infection in the neonate. The American journal of medicine, 70(2):417–422, 1981.

[6] M. E. Falagas and D. E. Karageorgopoulos. Extended-spectrum β-lactamase-producing organisms. Journal of Hospital infection, 73(4):345–354, 2009.

[7] Evelina Tacconelli. Antimicrobial use: risk driver of multidrug resistant microorganisms in healthcare settings. Current opinion in infectious diseases, 22(4):352–358, 2009.

[8] Donald A. Goldmann, Jeanne Leclair, and Ann Macone. Bacterial colonization of neonates admitted to an intensive care environment. The Journal of pediatrics, 93(2):288–293, 1978.

[9] Ebbing Lautenbach, Jean Baldus Patel, Warren B. Bilker, Paul H. Edelstein, and Neil O. Fishman. Extended-spectrum β-lactamase-producing escherichia coli and klebsiella pneumoniae: risk factors for infection and impact of resistance on outcomes. Clinical Infectious Diseases, 32(8):1162–1171, 2001.

[10] Janet Pasricha, Thibaud Koessler, Stephan Harbarth, Jacques Schrenzel, Véronique Camus, Gilles Cohen, Arnaud Perrier, Didier Pittet, and Anne Iten. Carriage of extended-spectrum beta-lactamase-producing enterobacteriacae among internal medicine patients in switzerland. Antimicrobial resistance and infection control, 2(1):20, 2013.

[11] Marc Lipsitch, Carl T. Bergstrom, and Bruce R. Levin. The epidemiology of antibiotic resistance in hospitals: paradoxes and prescriptions. Proceedings of the National Academy of Sciences, 97(4):1938–1943, 2000.

[12] B. S. Cooper, G. F. Medley, S. P. Stone, C. C. Kibbler, B. D. Cookson, J. A. Roberts, G. Duckworth, R. Lai, and S. Ebrahim. Methicillin-resistant staphylococcus aureus in hospitals and the community: stealth dynamics and control catastrophes. Proceedings of the National Academy of Sciences, 101(27):10223–10228, 2004.

[13] Hajo Grundmann and B. Hellriegel. Mathematical modelling: a tool for hospital infection control. The Lancet infectious diseases, 6(1):39–45, 2006.

[14] Ben S. Cooper. Confronting models with data. Journal of Hospital Infection, 65:88–92, 2007.

[15] Marie Forrester and Anthony N. Pettitt. Use of stochastic epidemic modeling to quantify transmission rates of colonization with methicillin-resistant staphylococcus aureus in an intensive care unit. Infection Control & Hospital Epidemiology, 26(7):598–606, 2005.

[16] Tat Ming Ng, Wendy X. Khong, Patrick N. A. Harris, Partha P. De, Angela Chow, Paul A. Tambyah, and David C. Lye. Empiric piperacillin-tazobactam versus carbapenems in the treatment of bacteraemia due to extended-spectrum beta-lactamase-producing enterobacteriaceae. PLoS One, 11(4):e0153696, 2016.

[17] Evelien de Jong, Jos A. van Oers, Albertus Beishuizen, Piet Vos, Wytze J. Vermeijden, Lenneke E. Haas, Bert G. Loef, Tom Dormans, Gertrude C. van Melsen, Yvette C. Kluiters, et al. Efficacy and safety of procalcitonin guidance in reducing the duration of antibiotic treatment in critically ill patients: a randomised, controlled, open-label trial. The Lancet Infectious Diseases, 16(7):819–827, 2016.

[18] Paul Turner, Sreymom Pol, Sona Soeng, Poda Sar, Leakhena Neou, Phal Chea, Nicholas P. J. Day, Ben S. Cooper, and Claudia Turner. High prevalence of antimicrobial-resistant gram-negative colonization in hospitalized cambodian infants. The Pediatric infectious disease journal, 35(8):856, 2016.

[19] Gabriel Birgand, Laurence Armand-Lefevre, Isabelle Lolom, Etienne Ruppe, Antoine Andremont, and Jean-Christophe Lucet. Duration of colonization by extended-spectrum β- lactamase-producing enterobacteriaceae after hospital discharge. American journal of infection control, 41(5):443–447, 2013.

[20] Ryan R. Wick, Louise M. Judd, Claire L. Gorrie, and Kathryn E. Holt. Unicycler: resolving bacterial genome assemblies from short and long sequencing reads. PLoS computational biology, 13(6):e1005595, 2017.

[21] Brian D. Ondov, Todd J. Treangen, Páll Melsted, Adam B. Mallonee, Nicholas H. Bergman, Sergey Koren, and Adam M. Phillippy. Mash: fast genome and metagenome distance estimation using minhash. Genome biology, 17(1):132, 2016.

[22] Kelly L. Wyres, Ryan R. Wick, Claire Gorrie, Adam Jenney, Rainer Follador, Nicholas R. Thomson, and Kathryn E. Holt. Identification of klebsiella capsule synthesis loci from whole genome data. Microbial genomics, 2(12), 2016.

[23] Richard McElreath. Statistical Rethinking: A Bayesian Course with Examples in R and Stan. CRC Press, 2018.

[24] Kathryn E. Holt, Heiman Wertheim, Ruth N. Zadoks, Stephen Baker, Chris A. Whitehouse, David Dance, Adam Jenney, Thomas R. Connor, Li Yang Hsu, Juliëtte Severin, et al. Genomic analysis of diversity, population structure, virulence, and antimicrobial resistance in klebsiella pneumoniae, an urgent threat to public health. Proceedings of the National Academy of Sciences, 112(27):E3574–E3581, 2015.

[25] Maria Bagattini, Valeria Crivaro, Anna Di Popolo, Fabrizio Gentile, Alda Scarcella, Maria Triassi, Paolo Villari, and Raffaele Zarrilli. Molecular epidemiology of extended-spectrum β-lactamase-producing klebsiella pneumoniae in a neonatal intensive care unit. Journal of Antimicrobial Chemotherapy, 57(5):979–982, 2006.

[26] Sean Conlan, Pamela J. Thomas, Clayton Deming, Morgan Park, Anna F. Lau, John P. Dekker, Evan S. Snitkin, Tyson A. Clark, Khai Luong, Yi Song, et al. Single-molecule sequencing to track plasmid diversity of hospital-associated carbapenemase-producing enterobacteriaceae. Science translational medicine, 6(254):254ra126–254ra126, 2014.

[27] Anna E. Sheppard, Nicole Stoesser, Daniel J. Wilson, Robert Sebra, Andrew Kasarskis, Luke W. Anson, Adam Giess, Louise J. Pankhurst, Alison Vaughan, Christopher J. Grim, et al. Nested russian doll-like genetic mobility drives rapid dissemination of the carbapenem resistance gene blakpc. Antimicrobial agents and chemotherapy, pages AAC–00464, 2016.

[28] Fredrik Bäckhed, Josefine Roswall, Yangqing Peng, Qiang Feng, Huijue Jia, Petia Kovatcheva-Datchary, Yin Li, Yan Xia, Hailiang Xie, Huanzi Zhong, et al. Dynamics and stabilization of the human gut microbiome during the first year of life. Cell host & microbe, 17(5):690–703, 2015.

[29] Amy Langdon, Nathan Crook, and Gautam Dantas. The effects of antibiotics on the microbiome throughout development and alternative approaches for therapeutic modulation. Genome medicine, 8(1):39, 2016.

[30] Jotham Suez, Niv Zmora, Gili Zilberman-Schapira, Uria Mor, Mally Dori-Bachash, Stavros Bashiardes, Maya Zur, Dana Regev-Lehavi, Rotem Ben-Zeev Brik, Sara Federici, et al. Post-antibiotic gut mucosal microbiome reconstitution is impaired by probiotics and improved by autologous fmt. Cell, 174(6):1406–1423, 2018.

